# Spatial receptive field structure of double-opponent cells in macaque V1

**DOI:** 10.1101/2020.01.20.913111

**Authors:** Abhishek De, Gregory D. Horwitz

**Author notes:** Correspondence: Gregory D. Horwitz, PhD, 1959 N.E. Pacific Street, Box 357330, HSB I-714, Seattle, WA, 98195.

## Abstract

The spatial processing of color is important for visual perception. Double-opponent (DO) cells likely contribute to this processing by virtue of their spatially opponent and cone-opponent receptive fields (RFs). However, the representation of visual features by DO cells in the primary visual cortex of primates is unclear because the spatial structure of their RFs has not been fully characterized. To fill this gap, we mapped the RFs of DO cells in awake macaques with colorful, dynamic white noise patterns. The spatial RF of each neuron was fitted with a Gabor function and a Difference of Gaussians (DoG) function. The Gabor function provided the more accurate description for most DO cells, a result that is incompatible with the traditionally assumed center-surround RF organization. A slightly modified (non-concentric) DoG function, in which the RFs have a circular center and a crescent-shaped surround, performed nearly as well as the Gabor model. For comparison, we also measured the RFs of simple cells. We found that the superiority of the Gabor fits over DoG fits was slightly more decisive for simple cells than for DO cells. The implications of these results on biological image processing and visual perception are discussed.

## INTRODUCTION

The spatial layout of chromatic signals influences color perception (Brown 1997; Monnier 2003; Singer 1994; Wachtler et al. 2001; Wandell 1993). A fundamental goal of visual neuroscience is to understand how the spatial processing of chromaticity by neurons mediate these effects. Double-opponent (DO) cells in primate primary visual cortex (V1) serve as a prime substrate for such an investigation because they encode colored edges (Conway 2001; Hubel and Wiesel 1968; Johnson et al. 2008; Livingstone and Hubel 1984). Neurons are defined as DO if they are cone-opponent (that is, if they receive antagonistic input from at least two types of cone photoreceptor in individual regions of their receptive fields (RFs)) and if they have opposite chromatic preferences in different parts of their RFs (Dow and Gouras 1973; Hubel and Wiesel 1968; Livingstone and Hubel 1984; Michael 1978; 1985; Poggio 1975; Thorell et al. 1984). These defining characteristics are undisputed, but the spatial structure of DO RFs is controversial.

DO cells were originally reported to have a center-surround RF organization (Hubel and Wiesel 1968; Livingstone and Hubel 1984; Michael 1978; 1985; Poggio 1975). More recent experiments have shown, however, that most DO cells are orientation-tuned, inconsistent with a center-surround RF organization (Johnson et al. 2008; 2001) but see (Conway 2001; Conway and Livingstone 2006). The differences in results between these studies are largely attributable to two factors. The first is the fact that different choices of visual stimuli can affect estimates of RF structure. Studies that reported center-surround RFs used uniform disks whereas those that reported orientation tuning used oriented gratings. A second factor is the difficulty inherent in inferring a 2-dimensional RF map from 1-dimensional data (e.g. disk size or grating orientation). At least some V1 cells have been shown to encode complex spatial patterns that cannot be predicted from orientation tuning measurements alone (Hegdé 2007; Tang 2018).

Knowing the complete 2-D spatial structure of DO cell RFs is important for three reasons. First, it is necessary for the construction of image-computable models, which can generalize to any image as input (Yamins and DiCarlo 2016). Second, it facilitates comparison with normative theories of image encoding (Caywood 2004; Hoyer 2000; Kellner 2013; Tailor 2000). Third, it elucidates the link between neurophysiology and perception. For example, center-surround DO RFs could contribute to differences in form perception defined by chromatic or luminance contrast (Gregory 1977; Livingstone 1987; Mullen 2002). Gabor-like RFs could contribute, additionally, to shape-from-shading—the ability to estimate the 3-D shapes of objects from shading cues (Kingdom 2003; Kunsberg 2018).

We stimulated V1 neurons with a white noise stimulus that varied independently in two spatial dimensions, as well as in color and time. We analyzed the data by spike-triggered averaging to identify DO cells and to measure their spatial RFs. We fit each spatial RF with two models that have been advanced previously to describe DO RFs: a difference of Gaussians (DoG) (Balasuriya 2003; Gao 2015; Lau 2008a; Lau 2008b; Spitzer 2005) and a Gabor function (Johnson et al. 2008; Yang 2013; Zhang 2012). We compared model fits between DO cells and simple cells—a benchmark cell type whose RF is well known to be better fit by a Gabor than a DoG function (Hubel and Wiesel 1968; Jones 1987; Moore IV 2012; Ringach 2002a; b).

The Gabor model outperformed the DoG model for most DO cells we studied. The goodness-of-fit of the Gabor model was similar for simple and DO cells. Some DO RFs consisted of a circular center and a crescent-shaped surround. A slight modification of the DoG model captured such RFs, performing nearly as well as the Gabor model for DO cells but poorly for simple cells. Together, these results show that simple and DO RFs are both well-described by the Gabor model, they are poorly described by the DoG model, and a center-crescent surround model is a reasonable description for many DO cells.

## METHODS

### General

All protocols conformed to the guidelines provided by the US National Institutes of Health and the University of Washington Animal Care and Use Committee. Data were collected from two male and two female rhesus macaques (Macaca mulatta) weighing 7–13 kg. Each monkey was surgically implanted with a titanium headpost and a recording chamber (Crist Instruments) over area V1. Eye position was continuously monitored using either an implanted monocular scleral search coil or an optical eye-tracking system (SMI iView X Hi-Speed Primate, SensoMotoric Instruments).

### Monitor calibration

Stimuli were presented on a cathode-ray tube (CRT) monitor (Dell Trinitron Ultrascan P991) with a refresh rate of 75 Hz against a uniform gray background y (x = 0.3, y = 0.3, Y = 43–83 cd/m^2^). Monitor calibration routines were adapted from those included in the Matlab Psychophysics Toolbox (Brainard 1997). Emission spectra and voltage-intensity relationships of each monitor phosphor were characterized using a spectroradiometer (PR650, PhotoResearch, Inc.). The color resolution of each channel was increased from 8 to 14 bits using a Bits++ video signal processor (Cambridge Research Systems, Ltd.) at the expense of spatial resolution; each pixel was twice as wide as it was tall.

### Task

Monkeys sat in a primate chair 0.7–1.0 m from a CRT monitor in a dark room during the experiments. The monkeys were trained to fixate a centrally located dot measuring 0.2 x 0.2° and to maintain their gaze within a square 1.0–2.0° fixation window. Successful fixation was rewarded, and fixation breaks aborted trials.

### Electrophysiological recordings

We recorded spike waveforms from well-isolated V1 neurons using extracellular tungsten microelectrodes (Frederick Haer, Inc.) that were lowered through dura mater by a hydraulic microdrive (Narishige, Inc. or Stoelting Co.). Electrical signals were amplified and digitized at 40 kHz online (Plexon, Inc.) and stored in a PC.

### Visual stimuli and experimental protocol

Each neuron was stimulated binocularly with white noise chromatic checkerboards (Horwitz et al. 2005; 2007). Each stimulus frame was a grid of 10 x 10 stimulus elements (stixels), and each stixel subtended 0.2 x 0.2°. The stimulus changed on every screen refresh. The intensity of each phosphor was modulated independently according to a Gaussian distribution with a standard deviation of 5–15% of the physically achievable range. The space-time averaged intensity of each phosphor was equal to its contribution to the background. Neuronal responses to the white noise stimuli were analyzed using spike triggered averaging (**Figure 1A**). Neurons that did not have clear spike-triggered averages (STAs) were passed over for data collection.

**Figure 1.**
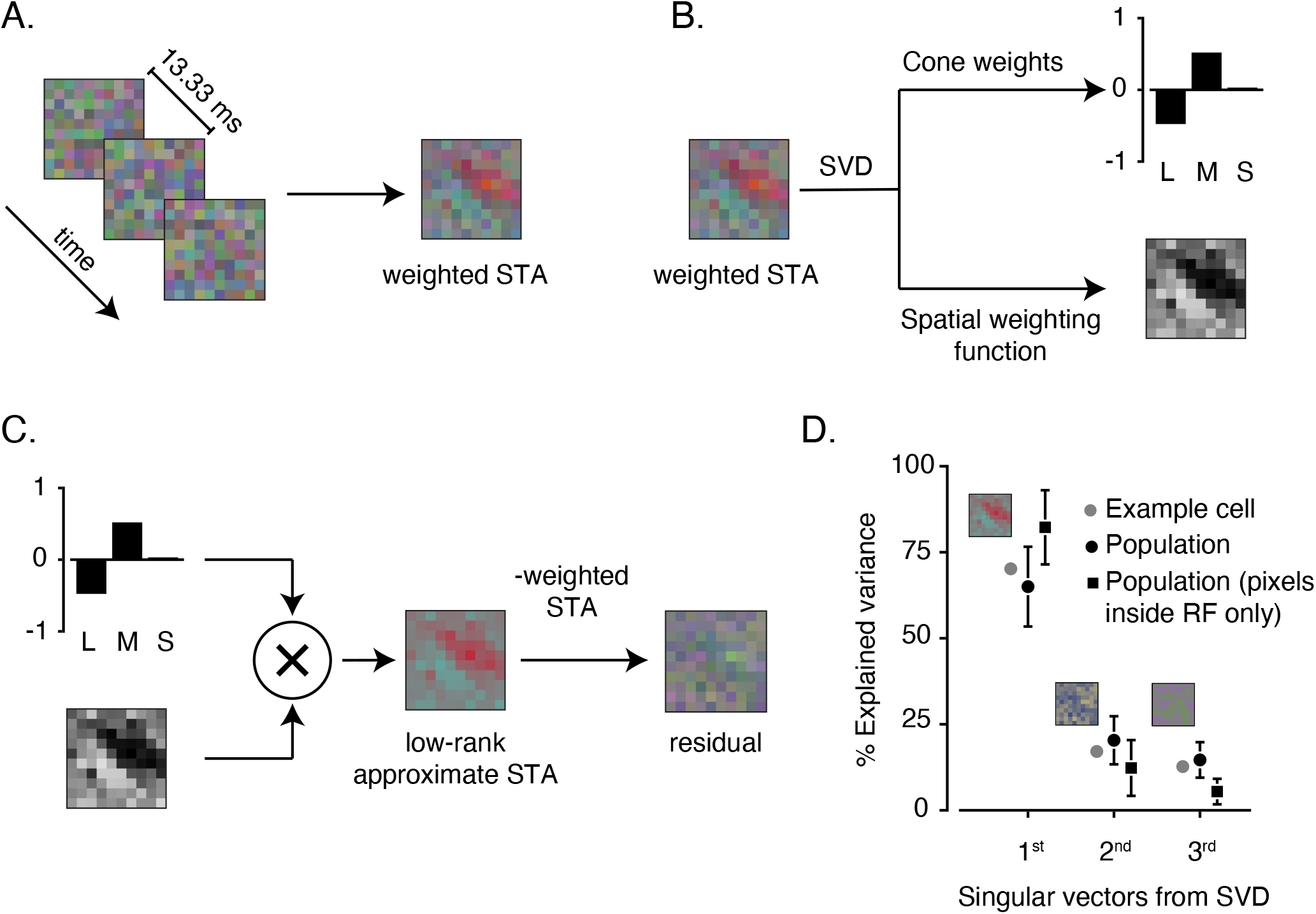
Derivation of cone weights and spatial weighting function. **A.** Calculation of the weighted STA (the weighted sum of the peak STA frame and two flanking frames) (right) from spike triggered white noise stimuli (left). **B.** Singular value decomposition (SVD) of the weighted STA reveals cone weights and spatial weighting function. **C.** Reconstructing a low-rank approximation of the weighted STA by multiplying cone weights and spatial weighting function. Subtracting the weighted STA from the the low-rank approximation yields the residual, which has little structure. **D.** Percent explained variance plotted against the three sets of singular vectors for the example cell and the population (mean ± SD). Cone weights and the spatial weighting function constitute the 1^st^ singular vectors. Percent explained variance was derived from the singular values using SVD over entire 10 stixels x 10 stixels of spatial weighting function (black circles) or omitting stixels outside of the RF (black squares).

### Cone weights and spatial RF

For each cell, we identified the frame from the STA that differed most from the background, based on the sum of squared red, green, and blue stixel intensities (negative intensities were defined as those below the contribution to the background). We then took the weighted average of the peak and the two flanking frames to create a 10 stixels x 10 stixels x 3 color channels tensor. The weight of each frame was proportional to the square root of sum of squared red, green and blue stixel intensities.

We reshaped the tensor into a 100 x 3 matrix, and used a singular value decomposition (SVD) to separate this weighted STA into a color weighting function and a spatial weighting function, defined as the first row and column singular vectors, respectively (**Figure 1B**) (Horwitz and Albright 2005). The color weighting function and the spatial weighting function captured most of the variance in the weighted STAs (**Figure 1C–D**).

The color weighting function, which quantifies neuronal sensitivity to modulations of the red, green, and blue phosphors of the display, was converted to cone weights that are assumed to act on cone contrast signals (Weller 2018). Cone weights were normalized such that the sum of their absolute values was 1 (Derrington et al. 1984; Horwitz and Albright 2005; Johnson et al. 2004). We analyzed only cells that were spatially opponent (**see Cell Screening**). As a result, each cell had cone weights with different signs in different RF subregions.

### Cell screening

We recorded from 393 V1 neurons and omitted 189 from the analyses on the basis of four criteria. Every neuron was required to have an STA with (1) high signal-to-noise ratio (SNR), (2) interpretable structure, (3) spatial opponency, and (4) cone weights that were either clearly opponent or clearly non-opponent. Below, we explain the rationale for each criterion and how it was implemented.

We excluded cells with low SNR because noisy STAs could lead to inaccurate estimates of color and spatial weighting functions. SNR was computed by comparing the peak STA frame to first STA frame and was defined as:

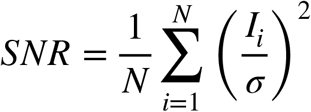

where *N* is the total number of elements within a frame: 10 stixels x 10 stixels x 3 color channels = 300 elements, *I* is the intensity of each element in the peak STA frame relative to the background, and *σ* is the standard deviation of the 300 elements that compose the first STA frame. The intensity of each element was divided by this standard deviation so that each element had (approximately) a standard normal distribution under the null hypothesis of no signal. We squared and summed these normalized intensity values and omitted from analysis the 60 cells for which this sum failed to reach a statistical threshold (p < 0.0001, *χ*^2^ test, df=300).

We excluded cells that combine cone inputs non-linearly because their STAs do not reflect their stimulus tuning accurately (Horwitz et al. 2005). We identified nonlinear neurons using a non-linearity index (NLI) (Horwitz et al. 2007). The NLI uses the STA and the spike-triggered covariance to find the maximally informative stimulus dimension under a multivariate Gaussian assumption (Pillow 2006). For each cell, we projected the stimuli shown in the experiment onto the maximally informative dimension and binned the projections, excluding the upper and lower 5% to avoid the influence of outliers. We calculated the average firing rate across the stimuli within each bin. The relationship between firing rate and stimulus projection was fit with three regression equations.

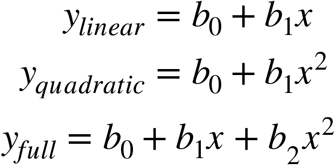

The goodness-of-fit of each regression was quantified with an *R*^2^ statistic. The NLI is defined as

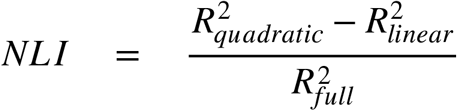

The NLI attains its theoretical maximal value of 1 when the inclusion of a linear term does not improve the regression fit. This would be the case, for example, for a V1 complex cell whose response is invariant to contrast polarity. NLI attains its theoretical minimum value of −1 when the inclusion of a quadratic term does not improve the regression fit as would be the case for a purely linear cell. Twenty-six cells were excluded on the basis that their NLI was > 0.

We excluded cells that were spatially non-opponent because these cells can be neither DO nor simple. We identified spatially non-opponent cells by analyzing the power spectrum of their spatial weighting functions. Spatially non-opponent cells, by definition, had maximal power in the lowest spatial frequency bin, which included power from 0 to approximately 0.7 cycles/°. This criterion excluded 54 cells. Other, stricter criteria excluded more cells but did not affect the main results.

We segregated simple cells from DO cells on the basis of cone weights, and we excluded neurons outside of these categories. Cells were classified as simple if their L-and M-cone weights had the same sign, accounted for 80% of the total cone weight, and individually accounted for at least 10%. Cells were classified as DO if they had large magnitude cone weights of opposite sign. DO_LM-opponent_ cells were defined as those that had L- and M-cone weights of opposite sign that together accounted for 80% and individually accounted for at least 20% of the total cone weight. DO_S-cone sensitive_ cells were cone-opponent and had an S-cone weight that accounted for at least 20% of the total. Forty-nine cells that were not categorized as simple, DO_LM-opponent_, or DO_S-cone sensitive_ were omitted from the analyses.

A total of 204 neurons contributed to the final pool (monkey 1: 42 simple, 57 DO_LM-opponent_, 37 DO_S-cone sensitive_; monkey 2: 11 simple, 11 DO_LM-opponent_, 1 DO_S-cone sensitive_; monkey 3: 18 simple, 14 DO_LM-opponent_, 6 DO_S-cone sensitive_; monkey 4: 1 simple, 4 DO_LM-opponent_, 2 DO_S-cone sensitive_).

### Model fitting of the spatial weighting function

We fit the spatial weighting function of each neuron with three models. Fitting was performed using the inbuilt MATLAB *fmincon* function to minimize the sum of squared errors between the spatial weighting function and the model fit. We describe each of the models below.

### Gabor model

The Gabor model was defined as:

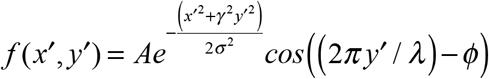

where (*x*′, *y*′) is obtained by translating the original coordinate frame to the RF center, (*x_c_, y_c_*), and rotating it by an angle *θ*.

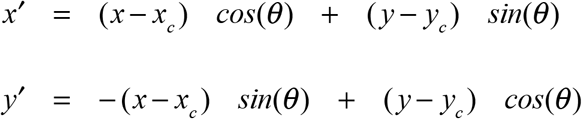

*λ* is the spatial period of the cosine component in °/cycle, and *ϕ* is the spatial phase. A spatial phase of *ϕ* = 0° produces an even-symmetric RF whereas spatial phase of *ϕ* = 90° produces an odd-symmetric RF. The two axes of the Gaussian envelope align with the *x*′ and the *y*′ axes. The parameter *A* is the amplitude, *γ* is the aspect ratio, and *σ* is the standard deviation of the Gaussian envelope along the *x*′ axis.

### Difference of Gaussians (DoG) model

The Difference of Gaussians (DoG) model can be written as:

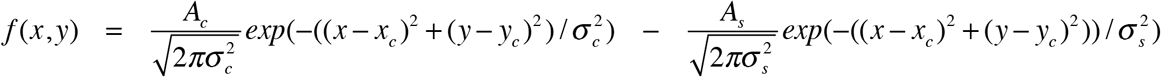

where *A_c_* and *A_s_* are the amplitudes of the center and surround. *σ_c_* and *σ_s_* are the standard deviations of the center and surround.

### Non-concentric DoG model

The non-concentric DoG model is identical to the DoG model but has two additional parameters (*x_s_, y_s_*) that allow the surround to be offset from the center (Dawis et al., 1984).

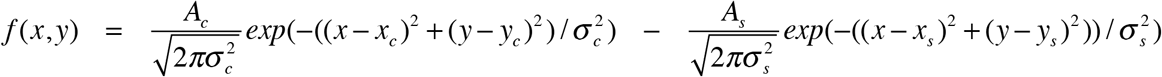

### Evaluating goodness of model fit: R

We evaluated the quality of model fits by calculating Pearson’s correlation coefficient (*R*) between the data and the model predictions. We used 5-fold cross validation, fitting the model with 80% of the data and testing the model on the remaining 20%. This procedure avoids overfitting, but it augments the natural bias of *R* towards zero because the training set is smaller than the actual data set. We report the averaged *R* across the 5 folds.

### Evaluating goodness of model fit: Fraction of variance unexplained

We evaluated the fraction of the variance unexplained from model fits. The fraction of unexplained variance was defined as the ratio of the residual sum of squared errors and the total sum of errors. For this calculation, we fit the models to the entire dataset for each neuron.

### Evaluating goodness of model fit: Bayesian information criterion

Model fits were further quantified using the Bayesian information criterion (*BIC*). Assuming that the model errors are independent and identically distributed according to a normal distribution, the *BIC* can be written as:

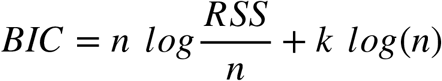

where *n* is the number of data points (*n* = 100), *RSS* is the residual sum of squared errors and *k* is the number of model parameters.

### Evaluating goodness of model fit: sum of squared errors

Model fits were further compared by calculating the sum of squared errors between the data and the fitted model. We used 5-fold cross-validation and report the averaged sum of squared errors across the 5 folds.

### Evaluating goodness of model fit: Prediction of spike-triggering stimuli

Model fits were further compared by calculating the ability of the models to predict spiking responses to white noise stimuli. Using 5-fold cross validation, the model was fit to 80% of the data and tested on the remaining 20%. Within the testing data, some segments of the white noise movie evoked a spike and most did not. We assessed the ability of the fitted model to classify movie segments that evoked a spike versus those that did not by projecting stimulus frames onto the 10 x 10 stixels (space) x 15 frames (time) spatial-temporal RF. The spatial-temporal RF was derived by combining the fitted spatial RF with the empirical time course of the STA. To compute classification performance, we constructed a receiver operating characteristic (ROC) from the spike and non-spike distributions of projections (Green and Swets 1966). We report the averaged area under the ROC across the 5 folds.

### Spatial opponency index

We defined a spatial opponency index (*SOI*) that quantifies the degree of antagonism across the RF as:

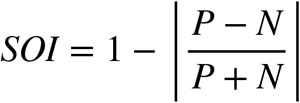

*P* was defined as the sum of positive values in the spatial weighting function. *N* was defined similarly but was the sum of negative values. If the sum of positive and negative values were matched, then *P* and *N* would be equal, and *SOI* would be equal to 1. On the contrary, if the RF consisted of a single subregion, then either *P* or *N* would equal 0, and so would the *SOI*.

## RESULTS

We analyzed the responses of 204 V1 neurons from 4 macaque monkeys that met our inclusion criteria (**see Methods**). RFs of neurons ranged in eccentricity from 1.7° to 8.4° (median = 4.7°).

### Cone weights

We classified neurons that met our inclusion criteria as simple cells or DO cells on the basis of spatial opponency and cone weights (**Figure 2**). Simple cells had large magnitude, L- and M-cone weights of the same sign that, together, accounted for 80% of the total cone weight (n=72). Neurons that were cone-opponent and spatially opponent were classified as DO cells. DO cells were further classified as LM-opponent (n=86) or S-cone sensitive (n=46) based on cone weight magnitudes and signs. Of the 46 DO_S-cone sensitive_ neurons recorded, 16 were S-(L+M), 26 were (S+M)-L, and 4 were (S+L)-M.

**Figure 2.**
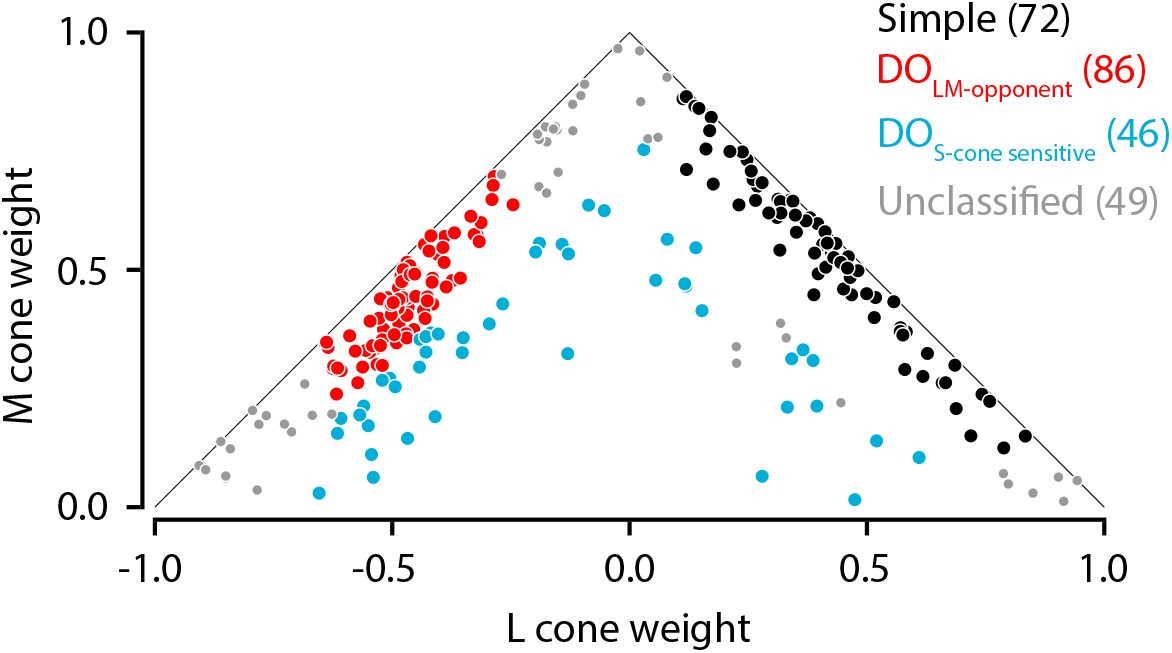
Normalized cone weights of simple (black), DO_LM-opponent_ (red), DO_S-cone sensitive_ (blue) and unclassified (gray) cells. The 49 “unclassified” cells survived all of the inclusion criteria except for the requirement of clear cone-opponency or non-opponency. M-cone weights were constrained to be positive. Points closer to the origin have larger S-cone weights than those far from the origin.

### Model comparison: Gabor vs. DoG

STAs of six example neurons illustrate patterns that we observed in the data (**Figure 3, 1^st^ row**). Statistical tests performed on individual phosphor channels, which are independent (**2^nd^–4^th^ rows**), reveal the color- and spatial-opponency of the four DO cells (C–F). Sensitivity to the three phosphors was converted to cone weights (**Figure 3, 5^th^ row**). Simple cell RFs consisted of adjacent ON and OFF regions (**Figure 3A & 3B**). Most simple cell RFs were elongated and clearly oriented (**Figure 3A**), but others were less so (**Figure 3B**). RFs of DO cells displayed similar features: some were clearly oriented (**Figures 3C & 3E**) whereas others had nearly circular RF centers and diffuse surrounds (**Figures 3D & 3F**).

**Figure 3.**
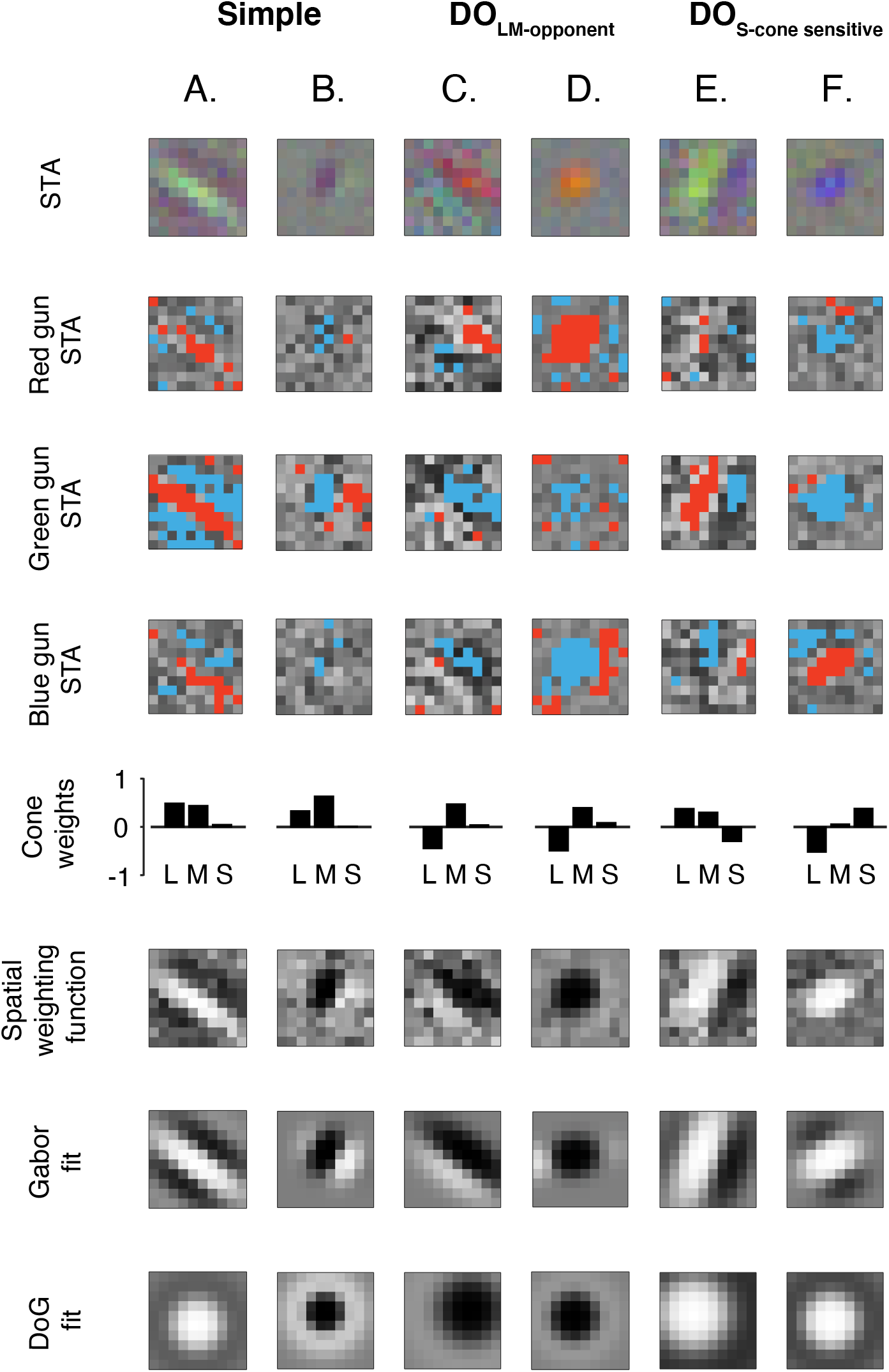
Gabor and Difference of Gaussians (DoG) model fits to spatial weighting functions of six example cells. Each spike-triggered average (top row) has been decomposed into red, green, and blue channel components. Significant stixels (z-test, p < 0.05) have been colored on the basis of their sign (red = positive, blue = negative). The quality of each model fit was quantified using cross-validated *R*. **A.** A simple cell with *R_Gabor_*= 0.77 and *R_DoG_*= 0.45 **B.** A simple cell with *R_Gabor_*= 0.67 and *R_DoG_*= 0.59 **C.** A DO_LM-opponent_ with *R_Gabor_*= 0.45 and *R_DoG_*= 0.30 **D.** A DO_LM-opponent_ cell with *R_Gabor_* = 0.90 and *R_DoG_*= 0.91 **E.** A DO_S-cone sensitive_ cell with *R_Gabor_*= 0.68 and *R_DoG_*= 0.44 **F.** A DO_S-cone sensitive_ cell with *R_Gabor_*= 0.78 and *R_DoG_*= 0.75.

To compare the spatial RF organization of simple and DO cells quantitatively, we converted the STAs to grayscale spatial weighting functions (**see Methods; Figure 3, 6^th^ row**) and fit them with a Gabor model (**Figure 3, 7^th^ row**) and a DoG model (**Figure 3, 8^th^ row**). Goodness-of-fit was quantified with cross-validated *R* between the data and the model predictions, a measure that allows fair comparison between models with different numbers of parameters (the Gabor has 8 parameters; the DoG has 6 parameters) but is biased towards zero for finite samples.

The Gabor model outperformed the DoG model for most of the cells tested (139/204, *R_Gabor_* > *R_DoG_*). The superiority of the Gabor model was consistent within each subgroup of cells: simple (p<0.001; Wilcoxon signed rank test; **Figure 4A**), DO_LM-opponent_ (p=0.07; **Figure 4B**) and DO_S-cone sensitive_ (p=0.006; **Figure 4C**). This result shows that DO cells, like simple cells, have RFs that are more accurately described by Gabor functions than DoG functions. However, the spatial RFs of simple and DO cells were not identical. The difference between *R_Gabor_* and *R_DoG_* was larger for simple cells than DO_LM-opponent_ or DO_S-cone sensitive_ cells (p<0.001 for each comparison; simple vs. DO_LM-opponent_; simple vs. DO_S-cone sensitive_ cells; Mann-Whitney U tests). The difference between *R_Gabor_* and *R_DoG_* was similar for DO_LM-opponent_ and DO_S-cone sensitive_ cells (p=0.25; Mann-Whitney U test). Analysis of fraction of variance unexplained by the model fits produced a similar result (**see Evaluating goodness of model fit: Fraction of variance unexplained**, Gabor model = 10.25±1.24%, DoG model = 15.63±0.97%).

**Figure 4.**
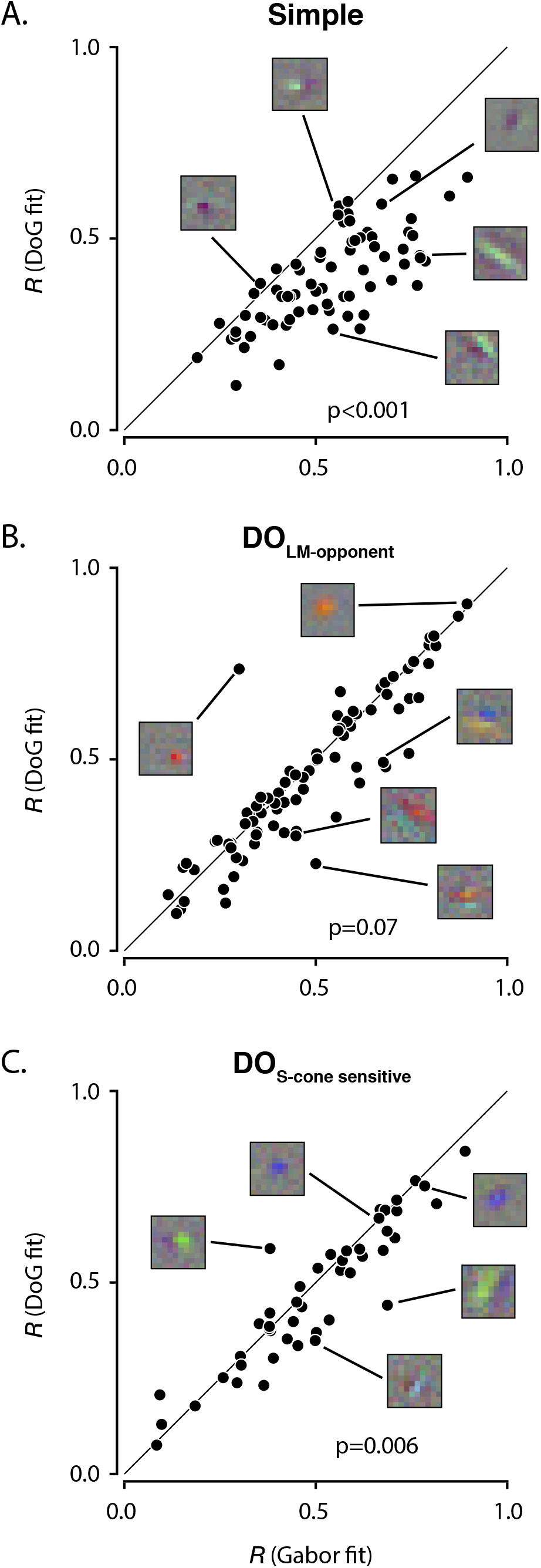
Comparison of Gabor and DoG model fits. Cross-validated *R* is plotted from Gabor fits and from DoG fits for simple (**A**), DO_LM-opponent_ (**B**), and DO_S-cone sensitive_ cells (**C**). Five example STAs are shown in each panel to illustrate the diversity of RF structures observed and their relationship to *R*.

We considered the possibility that systematic differences in SNR between DO cell STAs and simple cell STAs affected the model fits. For example, a spatial weighting function with low SNR would be equally well fit by a Gabor function as a DoG function even if the true RF organization was a DoG function. We therefore investigated the relationship between *R* and the SNR of the peak STA frame for each category of neurons (**see Methods** *for the definition of SNR*). As SNR increased, so did the goodness-of-fit of the Gabor model, which was similar across the three cell types (p=0.14, Kruskal-Wallis test; **Figure 5A**). This result shows that much of the error in the model fits is due to noise in the STAs, not to systematic errors in the Gabor model fits.

**Figure 5.**
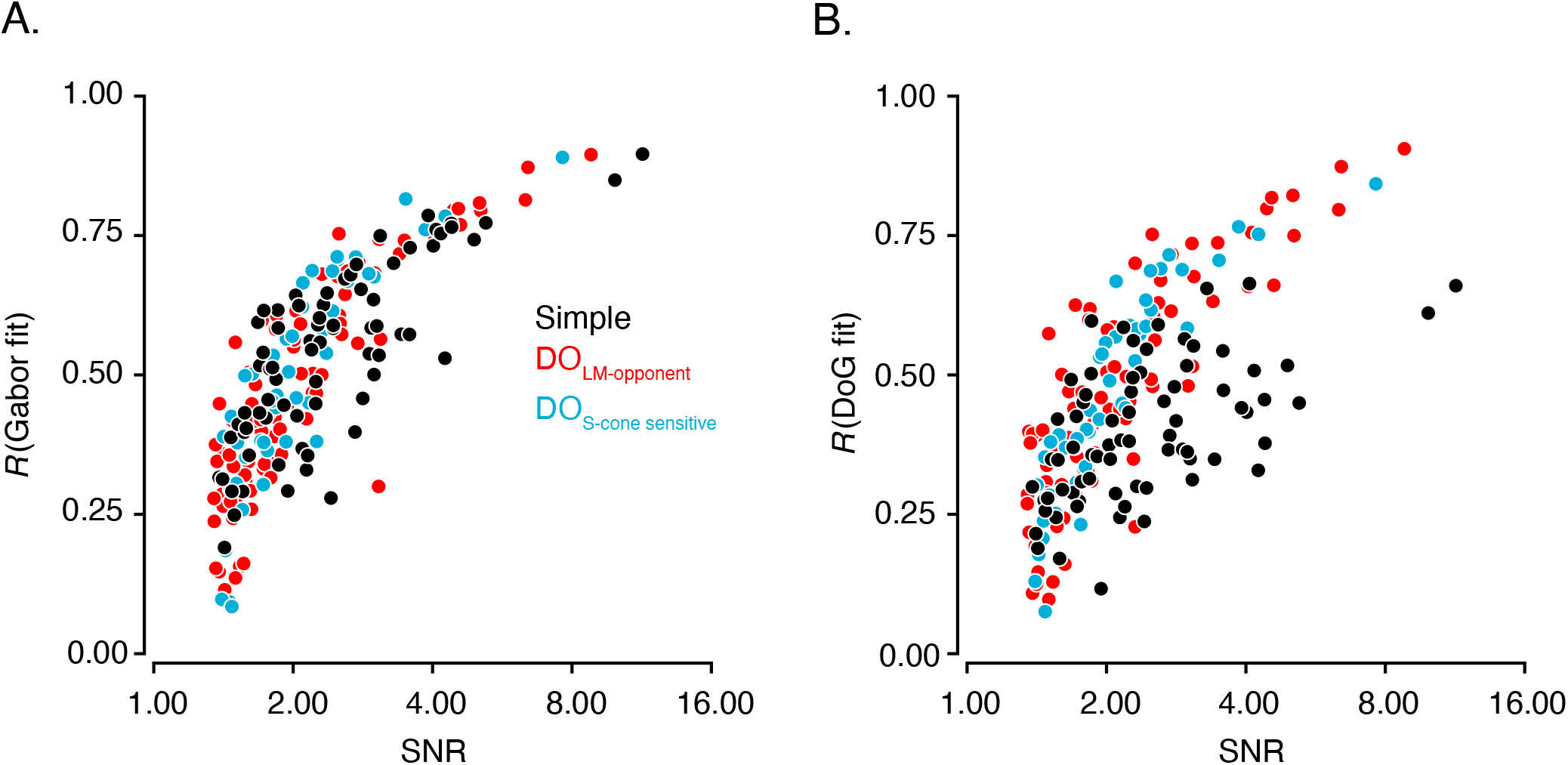
Analyses of Gabor and DoG model fits. **A.** Scatterplot of cross-validated *R* of Gabor fits vs. signal-to-noise ratios (SNR) of peak STA frames for simple cells (black), DO_LM-opponent_ cells (red) and DO_S-cone sensitive_ cells (blue). **B.** Identical to **A** but plotted for DoG fits.

A different result was obtained when SNR was compared to the goodness-of-fit of the DoG model. *R_DoG_* was lower for simple cells than for DO cells (**Figure 5B**, median for simple cells 0.38 vs. median for DO_LM-opponent_ 0.44 vs. median for DO_S-cone sensitive_ cells 0.44, p=0.07; Kruskal-Wallis test). This difference is clearest for cells with high SNR (p<0.0001, Kruskal-Wallis test on *R_DoG_* values for cells with SNRs above the median). A linear regression also confirmed that the relationship between (Fisher’s Z-transformed) *R_DoG_* and log_10_(SNR) differed across cell types (F-test, p < 0.0001)

To dissect the differences between simple cell and DO cell RFs more finely, we asked whether simple cell RFs are more frequently odd-symmetric or more elongated than those of DO cells. Either of these properties could degrade the quality of the DoG model fits relative to Gabor fits because DoG fits are constrained to be radially-symmetric. First, we analyzed the spatial phase of the best-fitting Gabor function, which makes the RF odd-symmetric, even-symmetric, or intermediate (*ϕ*, **see Methods**). Most simple cells were odd-symmetric (**Figure 6A**; mean = 57.2°), as were most DO_LM-opponent_ (**Figure 6B**; mean = 52.0°) and DO_S-cone sensitive_ cells (**Figure 6C**; mean = 52.0°).

**Figure 6.**
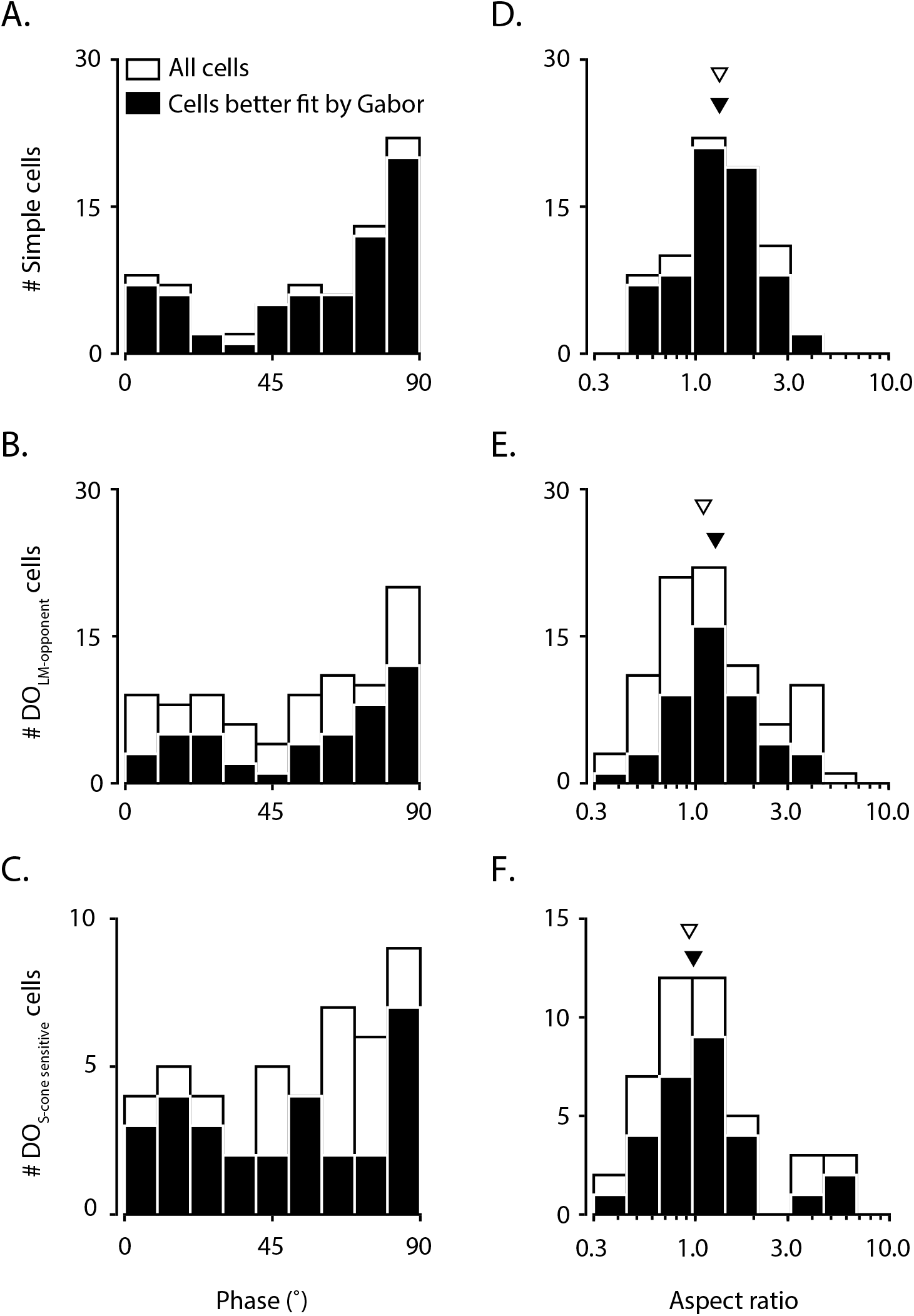
Analyses of Gabor model parameters for all cells (white) and cells that are better fit by the Gabor model than the DoG model (black). **A.** Best fitting phase (ϕ) of Gabor fits to simple cell spatial weighting functions. **B & C** Identical to **A** but for DO_LM-opponent_ cells and DO_S-cone sensitive_ cells, respectively. **D.** Best fitting aspect ratio (γ) of Gabor fits to simple cell spatial weighting functions. The median is plotted for all simple cell RFs (open triangle) and also for those that were better fit by the Gabor model (closed triangle). **E & F** Identical to **D** but for DO_LM-opponent_ cells and DO_S-cone sensitive_ cells, respectively.

Secondly, we analyzed the aspect ratio, which determines how elongated an RF is (*γ*, **see Methods**). Aspect ratios were larger for simple cells (**Figure 6D**; median = 1.33) than for DO_LM-opponent_ (**Figure 6E**; median = 1.09) or DO_S-cone sensitive_ cells (**Figure 6F**; median = 0.93). The difference in aspect ratio was statistically significant when all cells were considered (p=0.05, Kruskal-Wallis test). Restricting analysis to cells that were better fit by a Gabor model (*R_Gabor_* > *R_DoG_*) agreed qualitatively with the above results (**Figure 6A–F, black histograms**).

### The non-concentric DoG model

Gabor and DoG models are classic descriptions of DO RFs, but a third model, a center, with a crescent-shaped surround, has also been proposed (Conway 2001; Conway and Livingstone 2006). We formalized this idea by modifying the DoG model to allow the center and surround Gaussians to be non-concentric (**Figure 7A**) (Dawis 1984).

**Figure 7.**
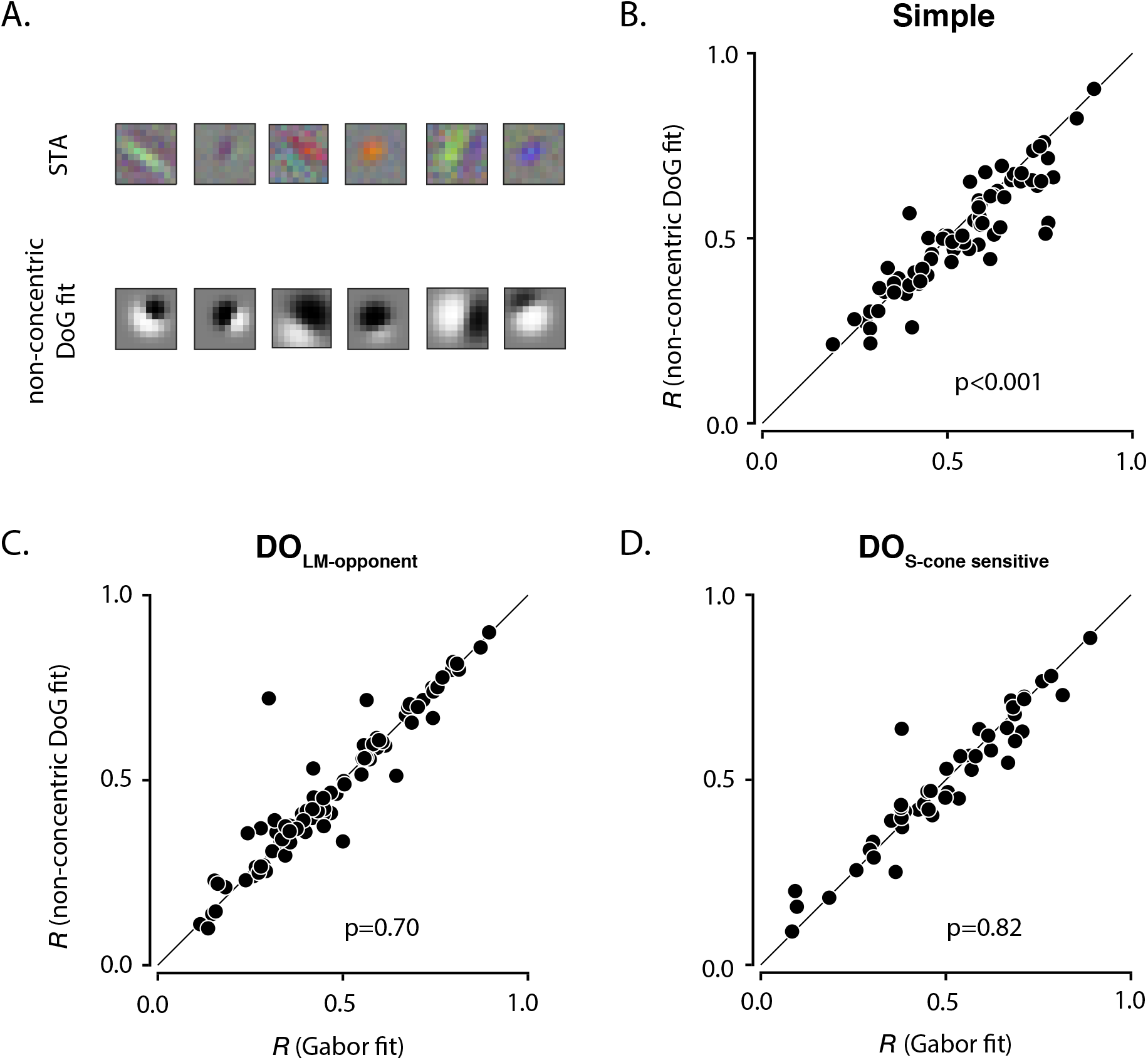
Comparison of non-concentric DoG and Gabor model fits. **A.** Non-concentric DoG fits to data from the six example cells in **Fig 3.** *R_non–concentric DoG_* = 0.54, 0.66, 0.41, 0.90, 0.60 and 0.78 from left to right. *R* from Gabor fits are plotted against *R* from non-concentric DoG fits for simple (**B**), DO_LM-opponent_ cells (**C**), and DO_S-cone sensitive_ cells (**D**).

We compared the quality of Gabor and non-concentric DoG fits for each cell. Simple cell RFs were better fit by the Gabor model (p<0.001; Wilcoxon signed rank test; **Figure 7B**) but DO_LM-opponent_ cells and DO_S-cone sensitive_ cell RFs were fit similarly by both models (p > 0.5; Wilcoxon signed rank tests; **Figure 7C and 7D**). The non-concentric DoG model is thus a reasonable description of DO cell RFs, but a Gabor model is superior for simple cell RFs.

### Analysis of spatial opponency

The antagonistic subfields of simple cell RFs in our data set were more nearly balanced than those of the DO cells. Spatial opponency indices (*SOI*s) were greater for simple cells (**Figure 8A**; median=0.91) than for DO_LM-opponent_ cells (**Figure 8B**; median=0.72) or DO_S-cone sensitive_ cells (**Figure 8C**; median=0.78) (p<0.001, Kruskal-Wallis test). As the *SOI* increased, so did the difference in goodness-of-fit of the Gabor model and the DoG model (r=0.39, p<0.001, Spearman’s correlation between *R_Gabor_* - *R_DoG_* and *SOI*; **Figure 8D**). This trend was not due to an increase in SNR (r=-0.08, p=0.25, Spearman’s correlation between SNR and *SOI*). The difference between *R_Gabor_* and *R_DoG_* was larger for simple cells than for DO cells even when analysis was restricted to the subset of cells with strong spatial opponency (p<0.001, Kruskal-Wallis test on *R_Gabor_* - *R_DoG_* values for cells with *SOI*s above the median). These results suggest that the superiority of the Gabor fits to simple cell RFs is not simply a consequence of their greater spatial opponency relative to DO cells.

**Figure 8.**
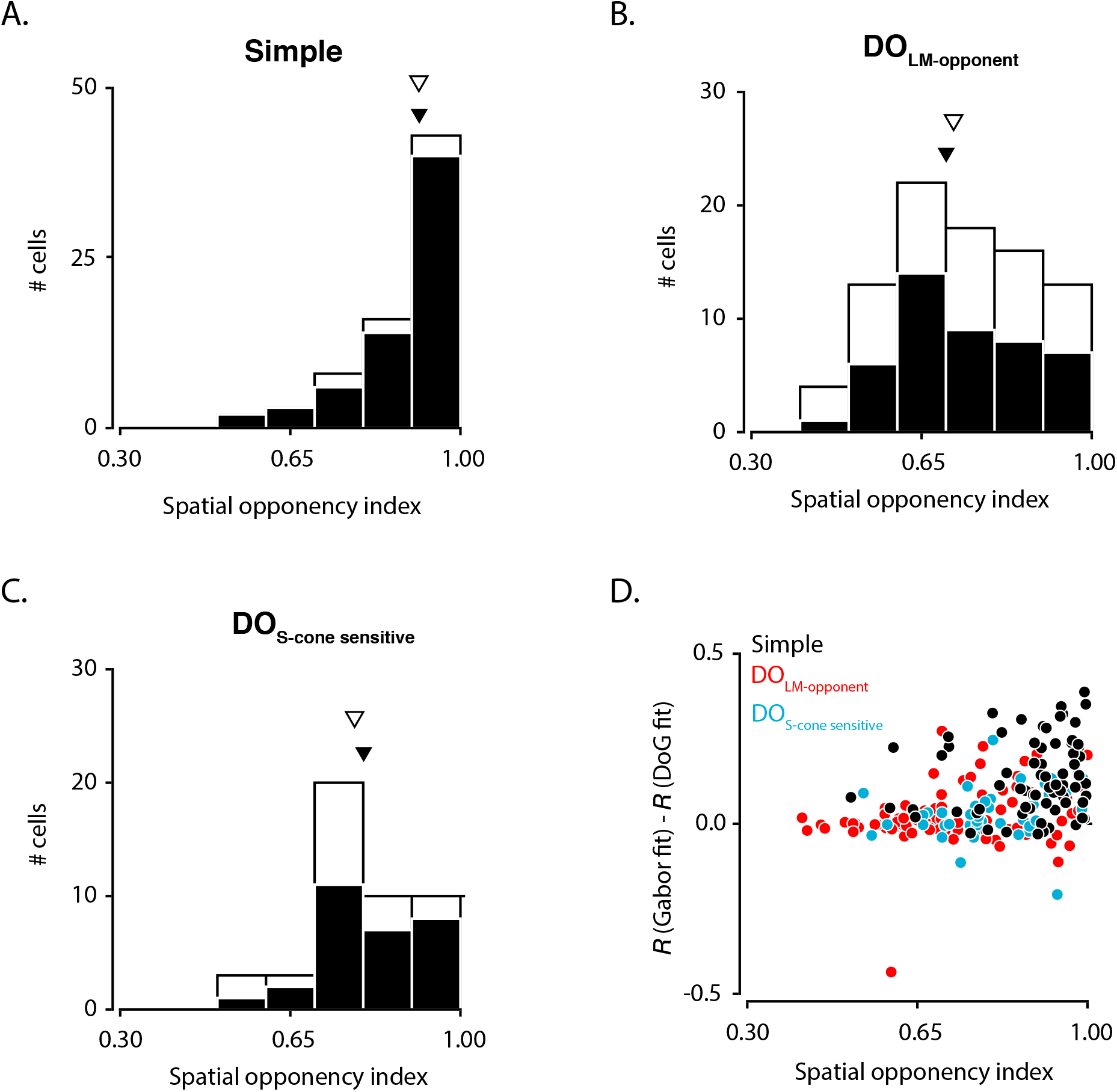
Analysis of spatial opponency. **A.** Histogram of spatial opponency indices (*SOI*s) for simple cells. The median *SOI* is plotted for all simple cell RFs (open triangle) and also of those that were better fit by the Gabor model (filled triangle). **B & C.** Identical to **A** but for DO_LM-opponent_ cells and DO_S-cone sensitive_ cells, respectively. **D.** Difference in *R* between Gabor and DoG fits is plotted against the *SOI* for simple (black), DO_LM-opponent_ (red) and DO_S-cone sensitive_ (blue) cells.

### Effect of eye movements

Eye movements cannot depend on which type of cell is recorded, but they could potentially favor one model over the other. To investigate whether this was the case in our data, we computed the median eye displacement from the average fixating eye position for each neuronal recording, and checked whether eye movements biased the model comparison results. The difference between *R_Gabor_* and *R_DoG_* was not significantly correlated with the magnitude of the eye displacement for any of the cell types (r=-0.08, p=0.49, Simple cells; r=-0.03, p=0.76, DO_LM-opponent_ cells; r=0.12, p=0.41, DO_S-cone sensitive_ cells; Spearman’s correlation between *R_Gabor_* - *R_DoG_* and median eye displacement; **Figure 9A**). Analysis of difference between *R_Gabor_* and *R_non-concentric DoG_* qualitatively agreed with the above result.

**Figure 9.**
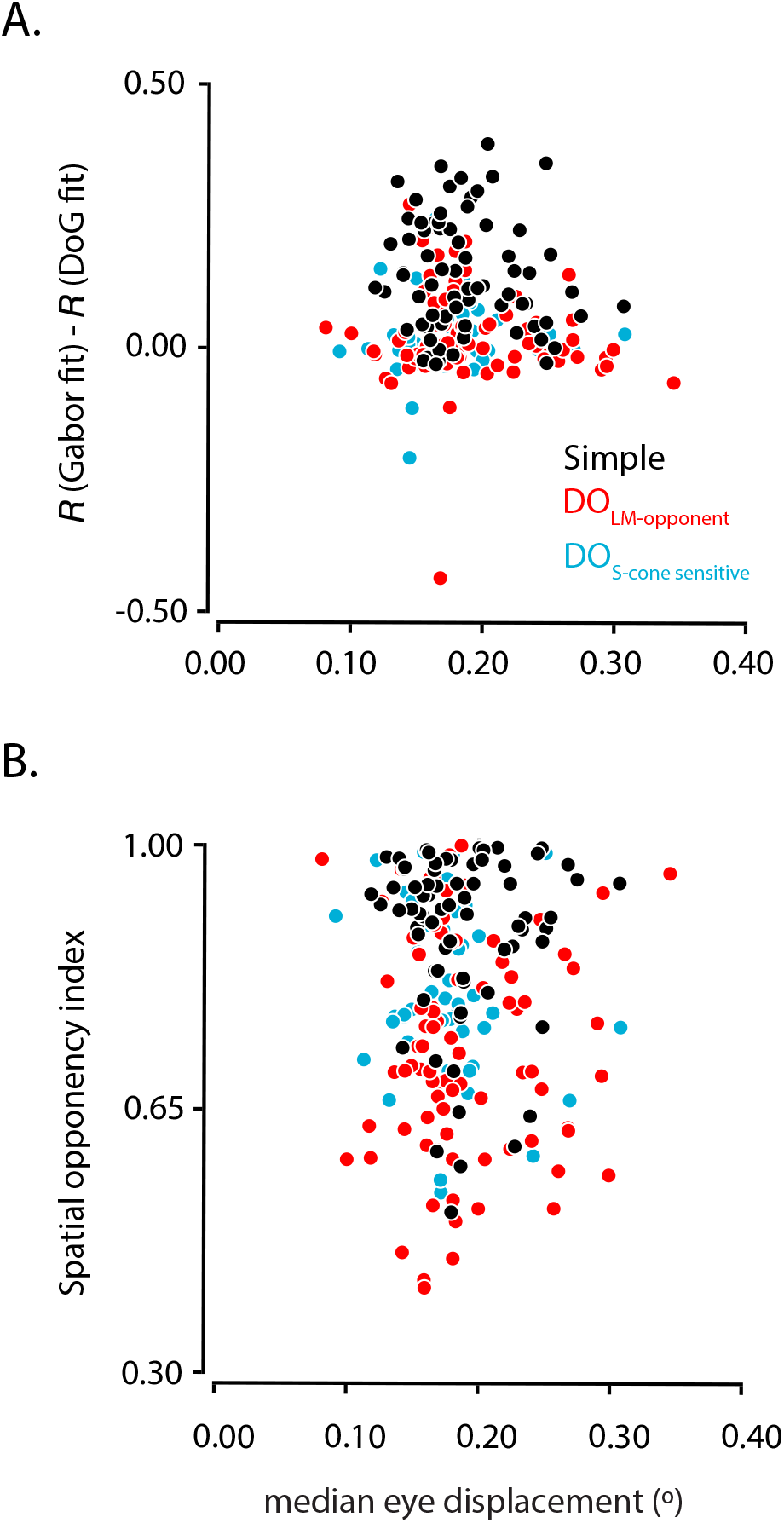
Effect of eye movements. **A.** Difference in *R* between Gabor and DoG fits is plotted against the median eye displacement for simple (black), DO_LM-opponent_ (red) and DO_S-cone sensitive_ (blue) cells. **B.** Spatial opponency index is plotted against the median eye displacement for simple (black), DO_LM-opponent_ (red) and DO_S-cone sensitive_ (blue) cells.

We also asked whether eye movements affected the spatial opponency we measured. We did not find any significant relationship between the *SOI* and the magnitude of eye movement (r=-0.06, p=0.59, simple cells; r=-0.02, p=0.80, DO_LM-opponent_ cells; r=-0.12, p=0.43, DO_S-cone sensitive_ cells; Spearman’s correlation between *SOI* and median eye displacement; **Figure 9B**). We conclude that our results on the spatial RFs of DO and simple cells are robust to eye movements.

## DISCUSSION

We measured the spatial RFs of macaque V1 DO and simple cells under identical conditions and compared them with rigorous statistical techniques. We report three new results. First, DO RFs, like simple cell RFs, were more accurately described by a Gabor model than a DoG model. Second, DO cells tend to have odd-symmetric RFs, similarly to simple cells. Third, DO RFs are more weakly spatially opponent than simple cell RFs. In summary, our results show that most DO cells lack a center-surround RF organization, the spatial RFs of DO and simple cells are broadly similar, and a center-crescent surround spatial structure describes DO cell RFs nearly as accurately as a Gabor function.

Below, we compare our results to those of previous studies and speculate on the neural wiring underlying simple and DO cells. We then discuss the robustness of our results to the statistics used to compare model fits. Finally, we discuss the potential roles of DO cells in image processing and how our findings have constrained these roles.

### Comparison with previous studies

Different studies have reached different conclusions about the spatial RF structure of DO cells in monkey V1 (Conway 2001; Conway and Livingstone 2006; Hubel and Wiesel 1968; Johnson et al. 2004; 2008; 2001; Livingstone and Hubel 1984; Michael 1978; Poggio 1975). Early investigations, mostly using circular spots of light, reported DO cells to have a concentric center-surround RF organization (Hubel and Wiesel 1968; Livingstone and Hubel 1984; Michael 1978; Poggio 1975). In some of these experiments, extremely large (>10° diameter) stimuli were tested, and these produced no response from DO cells. Such stimuli presumably recruit suppressive mechanisms from beyond the classical RF, which reconciles the lack of response to large, uniform stimuli with our observation that many DO RFs had imbalanced subfields. Later investigations using sparse noise stimuli measured 2-D RF structure and found that DO cell RFs have circular centers and crescent-shaped surrounds (Conway 2001; Conway and Livingstone 2006). Parallel investigations using drifting and rapidly flashed sinusoidal gratings concluded that DO cells have Gabor-like RFs (Johnson et al. 2004; 2008; 2001).

The lack of consensus about DO cell RF structure may reflect biases produced by different stimulus sets, incomplete RF descriptions, and different definitions of double-opponency. Sparse noise stimuli have the advantage of stimulating different parts of the RF independently and thus make no assumptions about the spatial structure of the RF (Conway 2001; Conway and Livingstone 2006). However, if every frame in a sparse noise stimulus consists of a pair of spots that are equal and opposite in contrast, then STAs, which are sums of these frames, will necessarily consist of equal parts contrast-increment and contrast-decrement, potentially producing an appearance of spatial opponency where none exists (Ben Lankow and Mark Goldman, personal communication).

Orientation tuning is compatible with a Gabor-like RF but does not necessarily make a strong case for it. In general, no 1-D measurement of spatial tuning completely constrains a 2-D RF profile. Knowing that a neuron is orientation-tuned is insufficient to simulate it in an image-computable model without making further assumptions.

Some studies included complex cells in the population of DO cells (Johnson et al. 2004; 2008; 2001). Complex cells do not abide by the classical definition of double-opponency because they do not have opposite color preferences in different parts of their RFs (Daw 1968; Dow and Gouras 1973; Hubel and Wiesel 1968; Livingstone and Hubel 1984; Michael 1978; 1985; Poggio 1975; Thorell et al. 1984). Some V1 neurons encode spatial phase and others do not. This distinction has proven to be useful in the achromatic domain, (e.g. for understanding the signal transformation between simple and complex cells) (Alonso 1998; Hubel 1962). Such a distinction is probably useful in the chromatic domain as well (Conway 2006).

Consistent with previous studies, we found that simple cell RFs are better fit by the Gabor model than by the DoG model and are usually odd-symmetric (Ringach 2002a; b). A novel contribution of the current study is the extension of this result to DO cells. The existence of odd-symmetric, chromatic edge detectors in the primate visual system was predicted on the basis of psychophysical experiments (Girard 1995; McIlhagga 2018).

### Are DO cells cone-opponent simple cells?

DO and simple cell RFs differ in detail but are similar in many ways. This similarity motivates the hypothesis that the primary difference between these two cell types is the sign of input they receive (indirectly) from the three cone photoreceptor classes. Indeed, the models proposed to underlie simple cell RFs can also be applied to some DO cells with only a minor change in the wiring (**Figure 10**).

**Figure 10.**
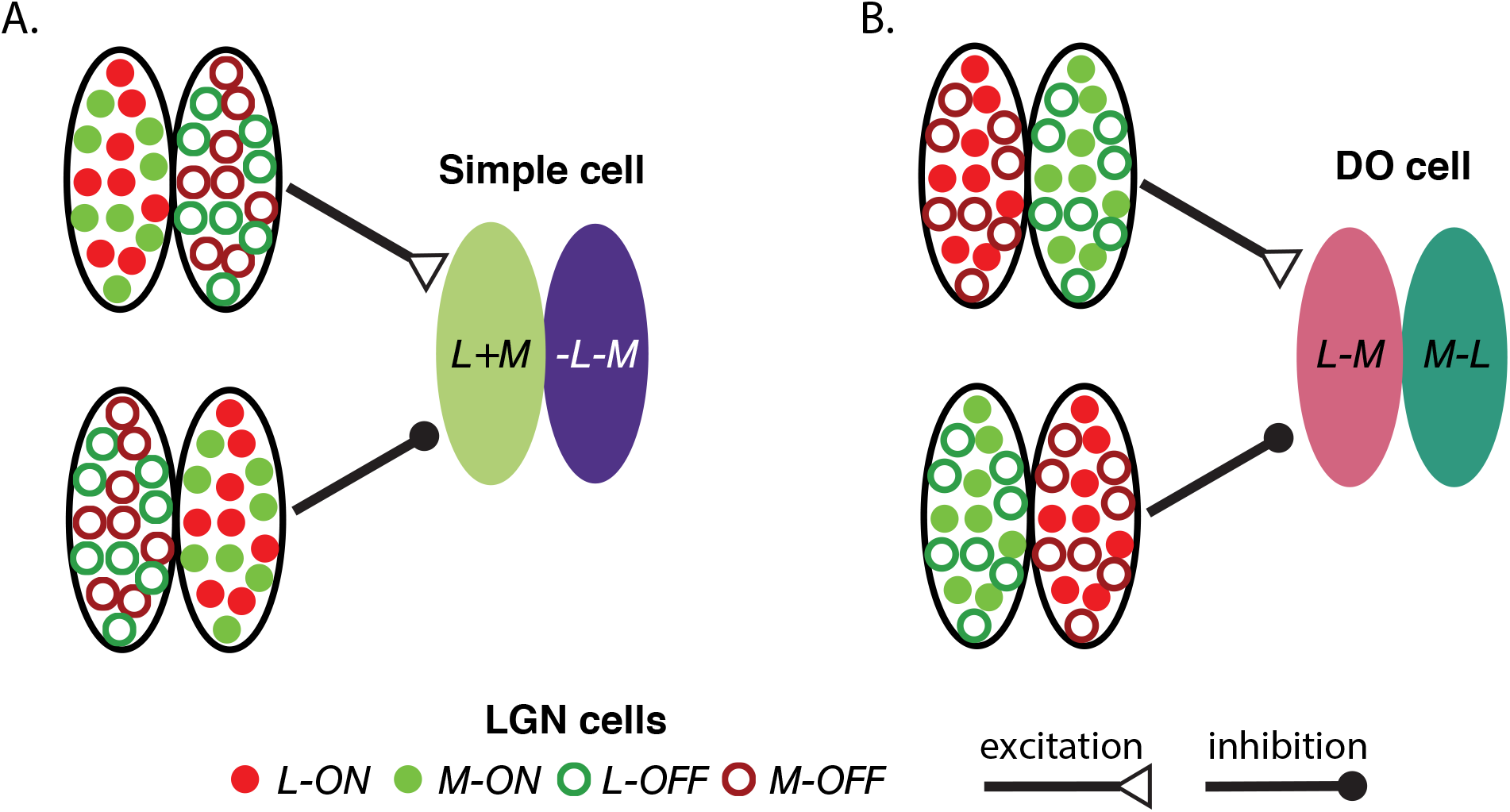
Schematic diagram of the circuitry proposed to underlie simple cell and DO cell RFs. **A.** A simple cell RF constructed from parvocellular LGN afferents. The ON subregion (L+M) is excited by L-ON and M-ON LGN cells and is inhibited by L-OFF and M-OFF LGN cells. Similarly, the OFF subregion (-L-M) is excited by L-OFF and M-OFF LGN cells and is inhibited by L-ON and M-ON LGN cells. **B.** Construction of a DO cell RF using the same set of parvocellular LGN cells that provide input to a simple cell. The L-M subregion is excited by L-ON and M-OFF LGN cells and is inhibited by L-OFF and M-ON LGN cells whereas the M-L subregion is excited by L-OFF and M-ON LGN cells and is inhibited by L-ON and M-OFF LGN cells.

A hallmark of simple cells is spatial linearity, a property mediated in part by push-pull excitation and inhibition (D. Ferster, 1988; D. Ferster, & Miller, K. D., 2000; Hirsch, 1998; Tolhurst, 1990). Some DO cells exhibit push-pull responses, consistent with the proposed similarly between them and simple cells (Conway 2006). However, whether the departures from linearity observed in DO cells exceeds expectations provided by the benchmark of simple cells is unclear. To answer this question, a useful next step is to compare quantitatively the degree of spatial linearity between DO and simple cells.

### Accuracy of RF structure and size

We carefully mapped V1 RFs using reverse correlation technique in awake macaque monkeys, similar to numerous previous studies (Conway 2001; Conway and Livingstone 2006; Horwitz and Albright 2005; Horwitz et al. 2005; 2007; Livingstone 1998; 2003; 1999; Pack 2006; 2003; Tsao 2003). Eye movements made by well trained monkeys during fixation blur measured RF maps, but this blurring would be expected to reduce or eliminate spatial structure, not to create it where is does not exist. All of the neurons we studied had spatially structured STAs. We analyzed the effects of eye movement on model fits and found no evidence that eye movements favored one model over the other (**Figure 9**). The distribution of eye positions was weakly anisotropic, showing more variance along the vertical than the horizontal axis, but the distribution of STA orientations did not exhibit a similar anisotropy. Additionally, these STA orientation preferences matched closely those measured directly with drifting sinusoidal gratings (data not shown), consistent with a previous study from our group (Horwitz et al. 2007), and inconsistent with the idea that eye movements produced artifactual, oriented STAs.

The RFs we measured were larger on average than those reported in anesthetized macaques at matched eccentricities (Hubel 1974; Van Essen et al. 1984). Eye movements surely contribute to this discrepancy as does the low contrast of the white noise stimulus and the large stixels in the white noise stimulus. Neurons with very small, spatially opponent RFs would be unlikely to respond to the stimulus and therefore would not have been studied. RF sizes measured using low contrast stimuli are approximately 2.3 times larger in area than those measured using high contrast stimuli (Sceniak et al. 1999). The effective contrast of our stimulus is low, owing to its Gaussian distribution and rapid refresh rate.

### Effects of cell categorization criteria

We distinguished simple cells from DO cells on the basis of cone weights. We applied a stricter criterion to L- and M-cone weights to categorize a cell as simple than as DO_LM-opponent_—a fact that is visible from the greater spread of L- and M-cone weights for simple cells than DO_LM-opponent_ cells (**Figure 2**). The rationale for this decision is the greater variability in estimated cone weights for non-opponent cells (Horwitz et al. 2007). Nevertheless, our results are robust to this decision (**Figures S1–2**).

### Alternative metrics for model comparison

We compared models using cross-validated correlation between data and model fits, but our results are robust to this choice. We repeated the model comparisons using the Bayesian Information Criterion, cross-validated sum of squared errors and crossvalidated prediction of spike-triggering stimuli. The results from all of these analyses agreed; RFs of DO and simple cells were more accurately described by a Gabor model than a DoG model (**Figure S3**), and the non-concentric DoG model provided a reasonable description of DO cell RFs but not simple cell RFs (**Figure S4**). We conclude that our conclusions are robust to the metric used to compare model fits.

### Role of DO cells in image processing

Our results show that some DO cells carry information about the phase and orientation of local chromatic variations. This information is useful for at least two visual computations. The first is shape-from-shading. Extraction of chromatic orientation flows in 2-D images is critical for accurate perception of 3-D shapes (Kingdom 2003; Kunsberg 2018; Zaidi 2006). In some displays, alignment of chromatic and luminance edges suppresses the percept of 3-D form whereas misalignment enhances the 3-D percept (Kingdom 2003). We speculate that signals from DO cells are integrated with those from simple cells to infer 3-D structure from 2-D retinal images. The similarity of RF structure between simple cells and DO cells may facilitate downstream integration of their responses. Second, DO cells might aid in inferring whether an edge in a visual scene is caused by the same material under different lighting conditions or by two different materials under the same lighting condition. An edge produced by a shadow falling across one half of a uniform material is a nearly pure intensity difference. On the contrary, an edge between two different materials under the same illumination is a consequence of spatially coincident intensity and spectral variations. A comparison of simple cell and DO cell responses could help to disambiguate material edges from illumination edges (Cavanagh 1991; Fine 2003; Olmos 2004; Tappen 2003).

## ACKNOWLEDGEMENTS

We thank Rich Pang and Fred Rieke for detailed comments on the manuscript. This work was funded by NIH EY018849 to Gregory D. Horwitz, NIH/ORIP grant P51OD010425, and NEI Center Core Grant for Vision Research P30 EY01730 to the University of Washington and R90 DA033461 (Training Program in Neural Computation and Engineering) to Abhishek De.

## SUPPLEMENTAL FIGURE LEGENDS

**Figure S1.**
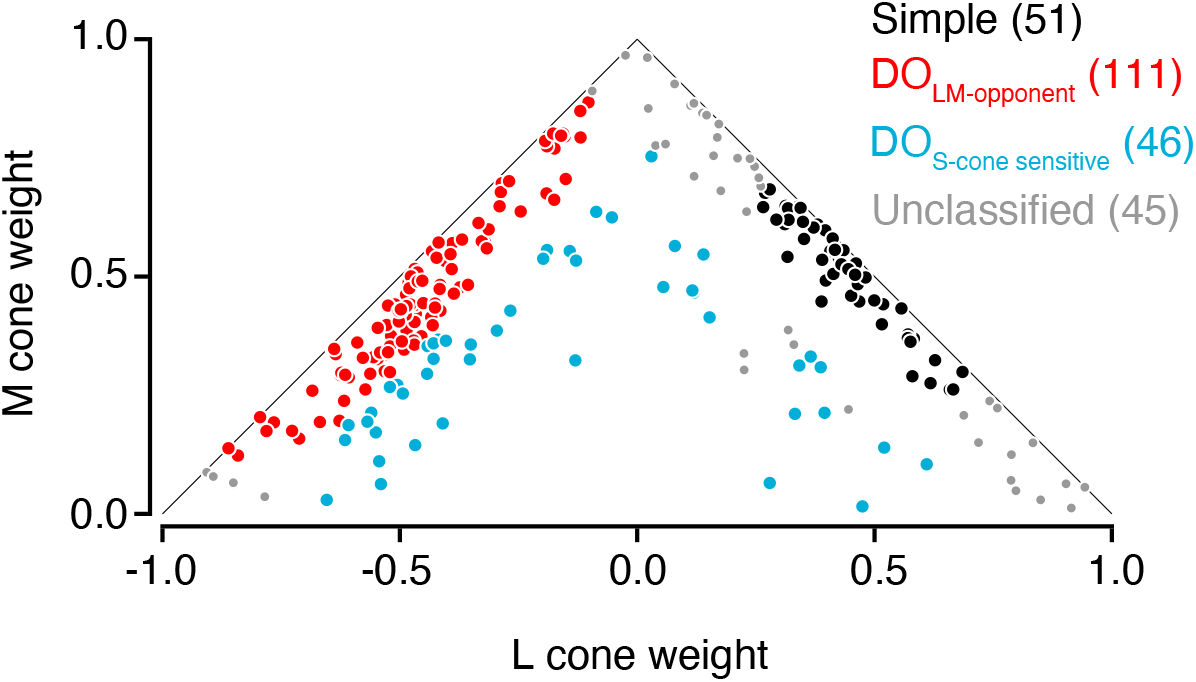
Reclassification of cells with reversed cone weight criteria. Shown are the normalized cone weights of simple (black), DO_LM-opponent_ (red), DO_S-cone sensitive_ (blue) and unclassified (gray) cells. Under these criteria, cells were classified as simple if the Land M-cone weights had the same sign, that together, accounted for 80% of the total cone weight and individually accounted for at least 20%. Cells were labeled as DO_LM-opponent_ if the L- and M-cone weights had opposite sign, together accounted for 80% and individually accounted for at least 10% of the total cone weight. Classification of DO_S-cone sensitive_ cells was unchanged from the description in the Methods.

**Figure S2.**
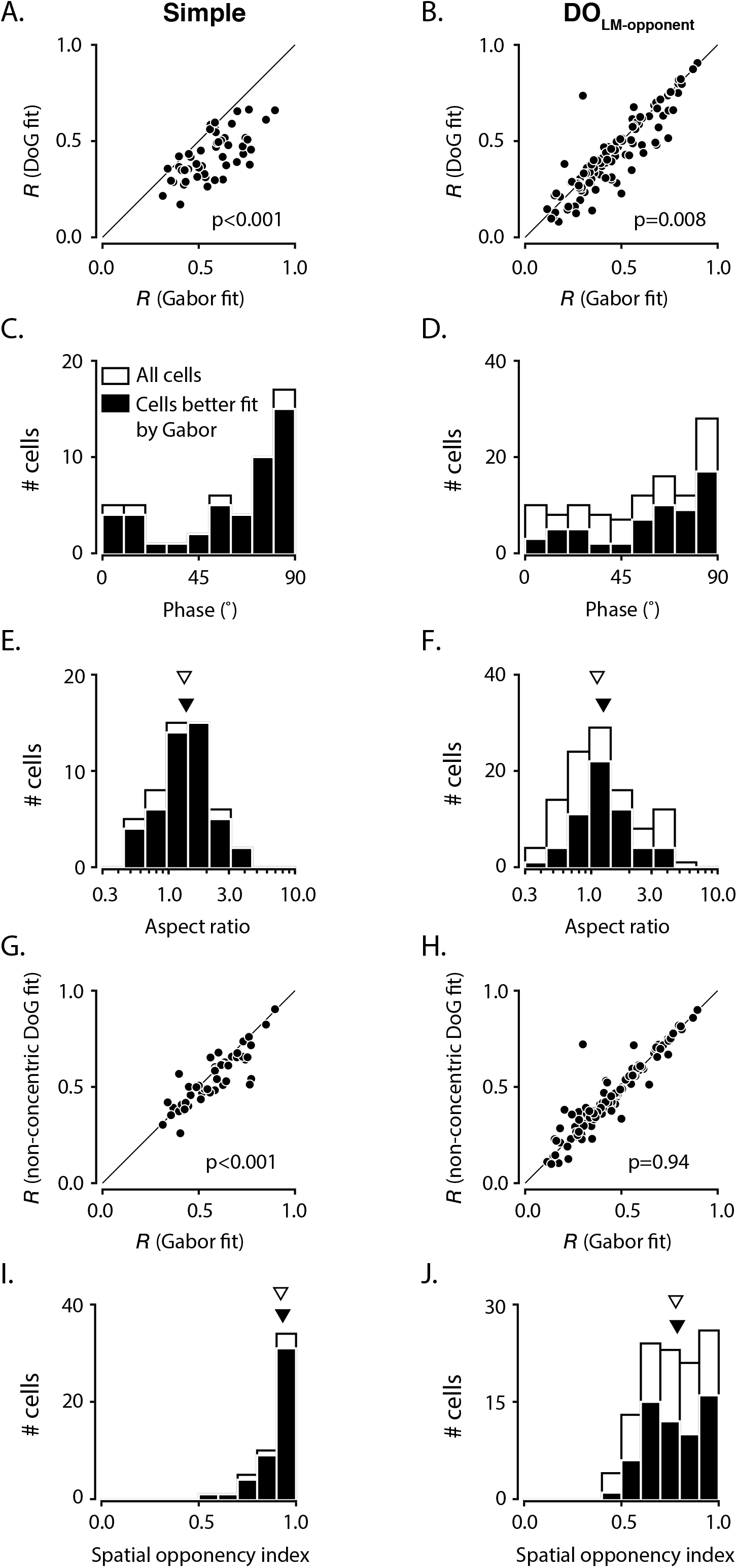
Model comparisons and spatial opponency analyses after reclassification of cells. **A.** Cross-validated *R* is plotted from Gabor fits and from DoG fits for simple cells. **B.** Identical to **A** but for DO_LM-opponent_ cells. **C.** Analysis of best fitting phase (ϕ) of Gabor fits to all simple RFs (white) and those that are better fit by the Gabor model than the DoG model (black). **D.** Identical to **C** but for DO_LM-opponent_ RFs. **E.** Analyses of best fitting aspect ratio (γ) of Gabor fits to all simple RFs (white) and those that are better fit by the Gabor model than the DoG model (black). The median γ is plotted for all simple cell RFs (open triangle) and also for cells better fit by Gabor model (closed triangle). **F.** Identical to **E** but for DO RFs. **G.** Cross-validated *R* is plotted from Gabor fits and from non-concentric DoG fits for simple cells. **H.** Identical to **G** but for DO_LM-opponent_ cells. **I.** Histogram of spatial opponency indices (*SOI*s) for simple cells based on spatial weighting functions. **J.** Identical to **I** but for DO_LM-opponent_ cells.

**Figure S3.**
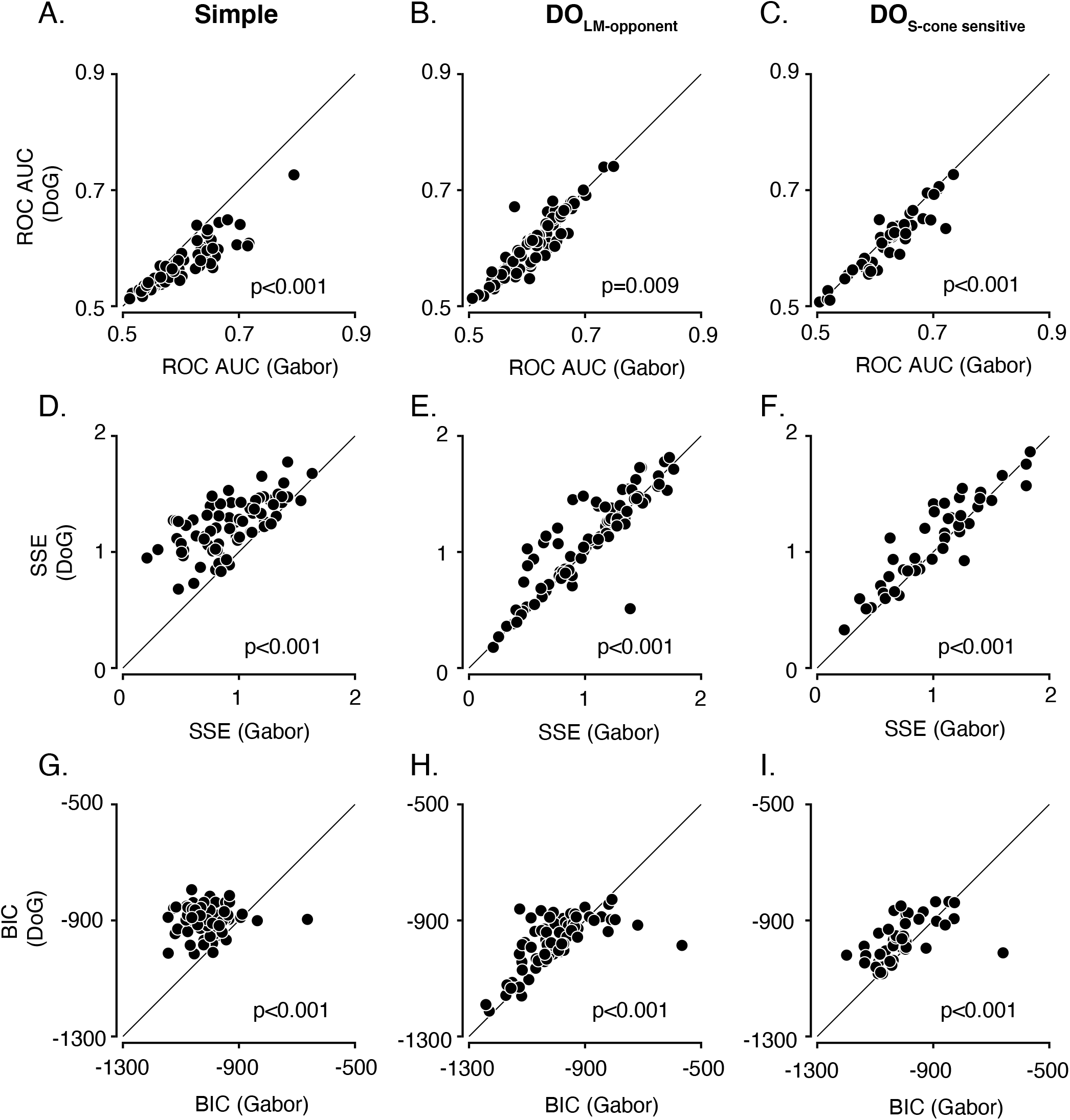
Comparison of Gabor and DoG model fits using three different analyses. **A.** Cross-validated prediction of spike-triggered stimuli using the area under receiver operating characteristics (ROC AUC) is plotted from Gabor fits and from DoG fits for simple cells. **B.** Identical to **A.** but for DO_LM-opponent_ cells. **C.** Identical to **A.** but for DO_S-cone sensitive_ cells. **D.** Cross-validated sum of squared errors (SSE) is plotted from Gabor fits and from DoG fits for simple cells. **E.** Identical to **D.** but for DO_LM-opponent_ cells. **F.** Identical to **D.** but for DO_S-cone sensitive_ cells. **G.** Bayesian Information Criterion (BIC) is plotted from Gabor fits and from DoG fits for simple cells. A better model fit yields a lower BIC. **H.** Identical to **G.** but for DO_LM-opponent_ cells. **I.** Identical to **G.** but for DO_S-cone sensitive_ cells.

**Figure S4.**
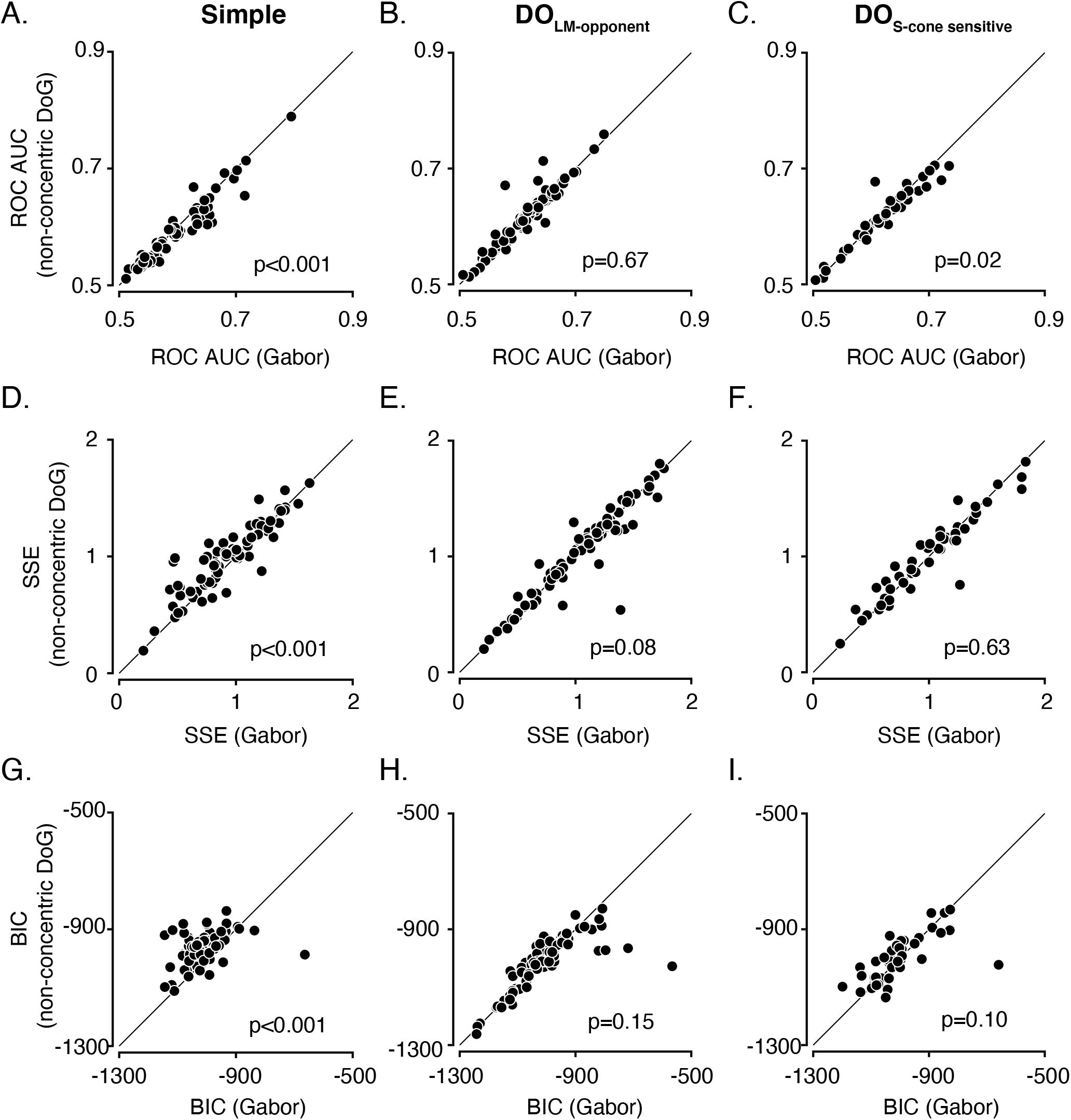
Comparison of Gabor and non-concentric DoG model fits using three different analyses. **A.** Cross-validated prediction of spike-triggered stimuli using the area under receiver operating characteristics (ROC AUC) is plotted from Gabor fits and from non-concentric DoG fits for simple cells. **B.** Identical to **A.** but for DO_LM-opponent_ cells. **C.** Identical to **A.** but for DO_S-cone sensitive_ cells. **D.** Cross-validated sum of squared errors (SSE) is plotted from Gabor fits and from non-concentric DoG fits for simple cells. **E.** Identical to **D.** but for DO_LM-opponent_ cells. **F.** Identical to **D.** but for DO_S-cone sensitive_ cells. **G.** Bayesian Information Criterion (BIC) is plotted from Gabor fits and from non-concentric DoG fits for simple cells. A better model fit yields a lower BIC. **H.** Identical to **G.** but for DO_LM-opponent_ cells. **I.** Identical to **G.** but for DO_S-cone sensitive_ cells.

